# FakeRotLib: expedient non-canonical amino acid parameterization in Rosetta

**DOI:** 10.1101/2025.02.27.640629

**Authors:** Eric W. Bell, Benjamin P. Brown, Jens Meiler

## Abstract

Non canonical amino acids (NCAAs) occupy an important place, both in natural biology and synthetic applications. However, modeling these amino acids still lies outside the capabilities of most deep learning methods due to sparse training datasets for this task. Instead, biophysical methods such as Rosetta can excel in modeling NCAAs. We discuss the various aspects of parameterizing a NCAA for use in Rosetta, identifying rotamer distribution modeling as one of the most impactful factors of NCAA parameterization on Rosetta performance. To this end, we also present FakeRotLib, a method which uses statistical fitting of small molecule conformer to create rotamer distributions. We find that FakeRotLib outperforms existing methods in a fraction of the time and is able to parameterize NCAA types previously unmodeled by Rosetta.

## Introduction

Within the past few years, protein structure modeling has experienced a revolution at the hands of deep learning-based technologies such as AlphaFold2,^1,2^ AlphaFold3,^3^ ESMFold,^4^ etc. However, the training of such models is dependent on the existence of plentiful structure data and the ability to represent the amino acid sequence with a finite vocabulary. This requirement can only be fulfilled for the twenty canonical amino acids, as modified and non-canonical residues are poorly represented in solved protein structures, and the amino acid alphabet used for sequence representation is restricted to canonical amino acid identities. These non-canonical amino acids (NCAAs) occupy an important place in natural biology, including post-translationally modified forms of canonical amino acids, stereochemically flipped “D-” amino acids, and various amino acid metabolites such as gamma-aminobutyric acid (GABA). In addition to natural biology, NCAAs have proved invaluable for peptide design^5,6^ (in particular peptide cyclization^7^), enzyme design,^8,9^ and probing individual residue functions in molecular biology experiments.^10^ Therefore, the inability to accurately model the conformations of these residues stands as a significant shortcoming of modern protein structure modeling.

Deep learning-based tools have begun to expand the amino acid space they are able to model with “all atom” modeling tools such as RoseTTAfold All-Atom^11^ and AlphaFold3^3^ making arbitrary chemistry comprehensible to their architectures. However, the performance of these tools leaves room to grow due to the insufficiency of available NCAA-containing structures in public databases. Therefore, to properly model proteins containing NCAAs, we must rely on more biophysical approaches to handle the wide diversity of chemistry which can occur for NCAAs. One approach is to collect the rotamers of a library of NCAAs before using them in downstream modeling tasks,^12–14^ but this approach incurs a high computational cost upfront and restricts the amino acid set to those which are explicitly represented in the library. Several studies have used molecular dynamics simulations to model the behavior of NCAAs in real time,^15–17^ but such methods are not well suited for tasks such as *in silico* peptide design which require the screening of hundreds of potential NCAAs at various positions along the sequence. Therefore, more coarse-grained methods such as Rosetta stand as the most appropriate tool for the modeling of NCAAs.

Rosetta first tackled NCAA modeling with MakeRotLib,^18^ an approach which calculated rotamer distributions through energetic minimization of sidechains using a hybrid Rosetta and CHARMM potential, but this method is severely limited by its runtime (days of walltime for the more flexible amino acids, even with MPI-based multithreading), its request for an initial guess of the position of rotamer distribution centroids, and its requirement that the torsions of the NCAA are found in its CHARMM-based energy repository. Since then, only a few methods have been published that address the rotamer generation problem for arbitrary NCAAs. One such method, AutoRotLib^5^ was recently published to expand the NCAA chemistry which could be parameterized by Rosetta, but it depends on proprietary OpenEye software and has a similarly long runtime compared to MakeRotLib. In this manuscript, we benchmark current methods for parameterization of NCAAs in Rosetta and discuss the impact of various aspects of NCAA parameterization on Rosetta’s modeling performance. In addition, we introduce our own method for NCAA rotamer parameterization, FakeRotLib, which uses open-source small molecule toolkits along with cartesian-space mixture models to efficiently create rotamer distributions.

## Results and Discussion

### Atom type and partial charge assignment

In order to test how atom typing impacts the performance of Rosetta in molecular modeling tasks, we created “non-canonical” forms of each of the canonical amino acids. In this test, we generated these “non-canonical” residues by replacing the “atom” block of the default amino acid parameter files with blocks that had been generated by three other methods: 1. Rosetta’s molfile_to_params_polymer.py script, 2. BCL-generated partial charges (summed signa and pi charges for each atom of the dipeptide)^19^ passed into molfile_to_params_polymer.py, and 3. default Rosetta atom types with partial charges drawn randomly from a [-1,1] uniform distribution (and subsequently corrected so that the sum of partial charges equaled the formal charge of the amino acid). The tasks we decided to use to benchmark Rosetta performance were a rotamer recovery task, in which all residues are mutated into their respective “non-canonical” forms and repacked to recover the native sidechain torsion angles, and a sequence recovery task, in which the native backbone is designed by Rosetta using the “non-canonical” amino acids with the aim of recovering the native amino acid sequence. On both rotamer and sequence recovery tasks, we have determined that Rosetta is robust with respect to partial charge values and minute differences in atom typing, as long as the partial charge values are reasonable (**Figure 1**). However, Rosetta is not completely numb to partial charges either, as randomizing partial charges leads to a decrease in modeling performance.

**Figure 1.**
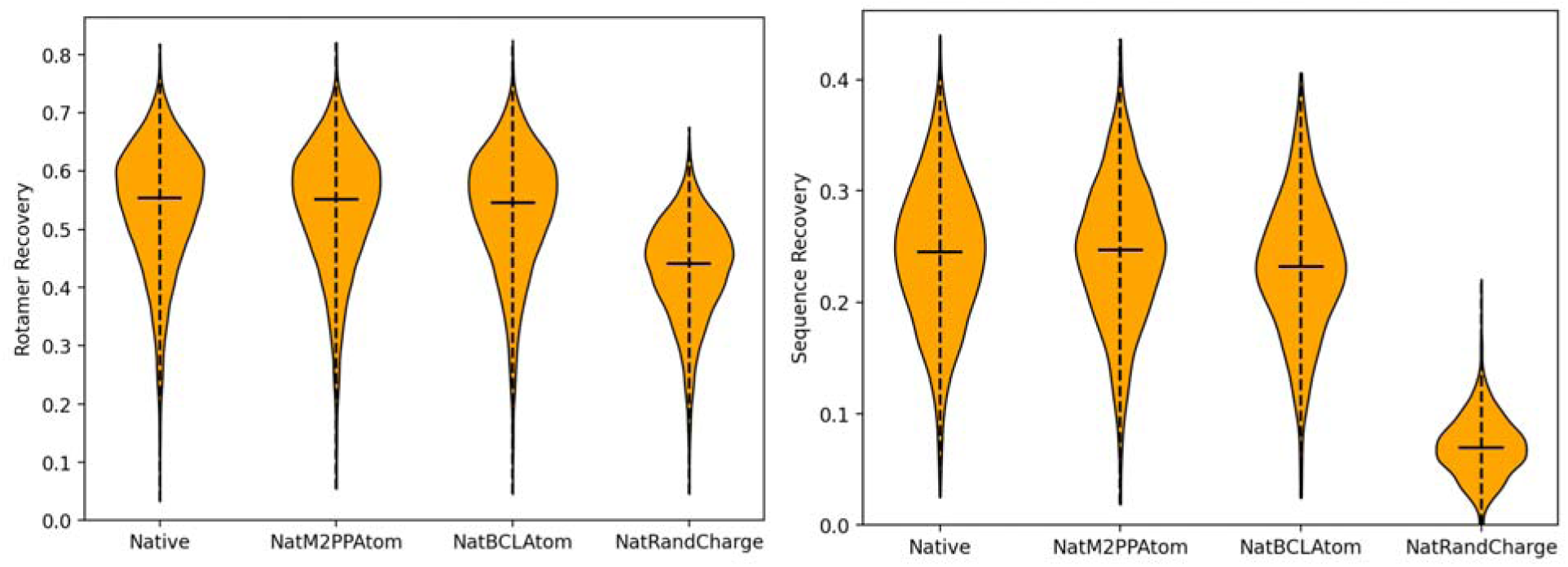
Rotamer recovery (left) and sequence recovery (right) performance of default Rosetta (Native), molfile_to_params_polymer.py assigned atoms (NatM2PPAtom), BCL partial charges (NatBCLAtom), and random partial charges (NatRandCharge).

### Sidechain geometry

Similar to how we tested the atom typing, we generated “non-canonical” forms of the canonical amino acids by replacing the internal coordinates block with internal coordinates of three different sources: 1. A geometry generated by the BCL conformer generator,^20^ 2. Molecular mechanics-optimized geometry via RDKit’s^21^ UFF^22^ and MMFF^23^ implementations, and 3. Quantum mechanically optimized geometry using Gaussian.^24^ We then assessed the performance of these geometries using the same tasks as before (**Figure 2**). In contrast to atom typing, Rosetta is sensitive with respect to molecular geometry, likely because the default movement set in Rosetta is restricted to torsional space, keeping bond lengths and angles fixed. However, clearly the bond lengths and angles are hyper-optimized with respect to the Rosetta scoring function. High-rigor quantum mechanical geometry was outperformed by more efficient molecular mechanical geometries in both rotamer and sequence recovery tasks, i.e., the rigor with which the geometry is created does not necessarily guarantee increases in performance. Therefore, an optimal methodology with respect to Rosetta performance would be one which exactly replicates the Rosetta-based geometry of the canonical amino acids *ab initio*; molecular mechanics-based optimization seems to offer the best approximation of this.

**Figure 2.**
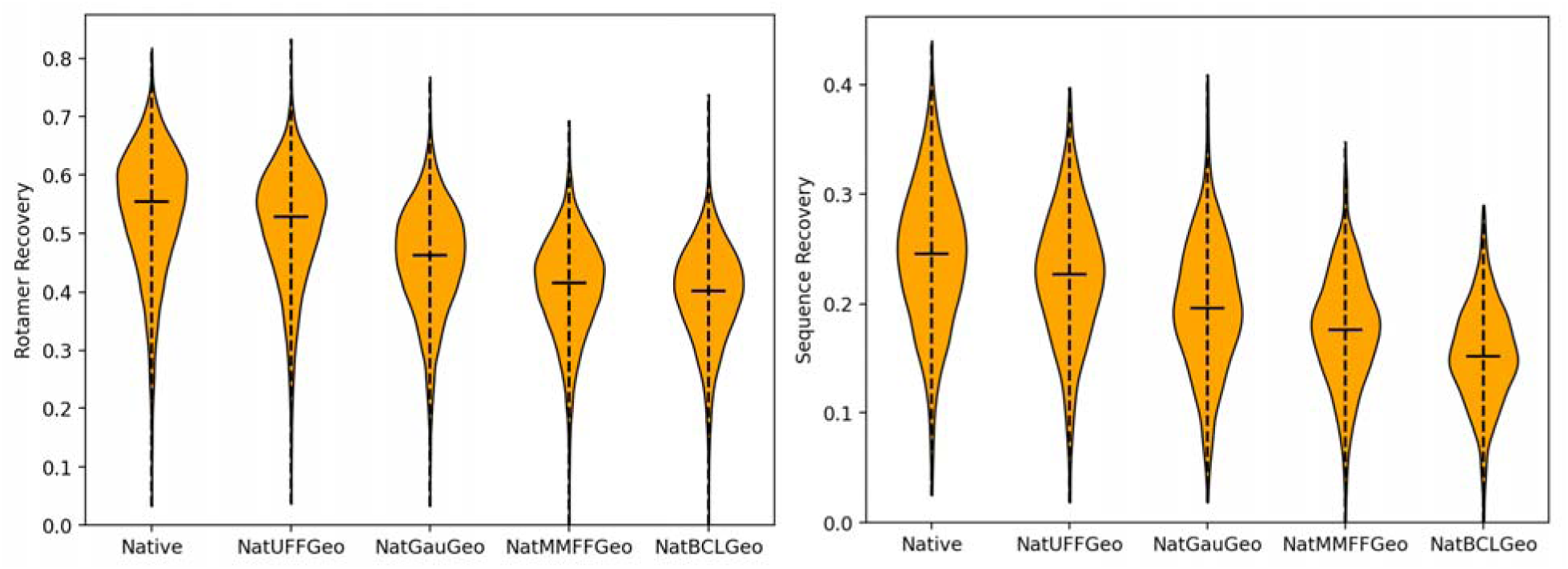
Rotamer recovery (left) and sequence recovery (right) performance of default Rosetta (Native), UFF-optimized geometry (NatUFFGeo), QM-optimized geometry (NatGauGeo), MMFF94-optimized geometry (NatMMFFGeo), and BCL-optimized geometry (NatBCLGeo).

### Sidechain Rotamers

Finally, we test different methods of generating rotamers for non-canonical amino acids and their impact on Rosetta performance. For these benchmarks, all except the “native” and “AutoRotLib” methods used the partial charges/atom types from molfile_to_params_polymer.py and geometry optimized by UFF. The purpose of doing this as opposed to using native geometries and atom types is to faithfully represent the performance one can expect to attain for a non-canonical amino acid without *a priori* knowledge. We tested seven unique methods of generating rotamers: 1. copying the rotamers from the native residues, a.k.a. using “parent” rotamers 2. MakeRotLib, a protocol developed by Renfrew et. al.^18^ which minimizes sidechain conformations via a hybrid Rosetta/CHARMM energy function, thus creating minimized rotamer wells 3. BCL, which generates poses of the sidechain via the BCL conformer generator^20^ and utilizes them as PDB rotamers 4. RDKit, which does the same as the previous method but using RDKit instead of BCL 5. FakeRotLib, an approach which creates a rotlib file similar to MakeRotLib by fitting the conformational distribution made by RDKit in cartesian space with a Bayesian Gaussian Mixture Model (BGMM) 6. AutoRotLib, an approach developed by Holden and Pavlovicz et. al.^5^ which uses OpenEye tools^25^ to mimic the MakeRotLib protocol 7. Providing no rotamer information to Rosetta and allowing it to model sidechains via uniform sampling (negative control).

From these results, we have determined that the “parent” rotamer approach performs the most closely to default Rosetta for rotamer recovery, followed by FakeRotLib, MakeRotLib, AutoRotLib, RDKit, BCL, and finally, no rotamers (**Figure 3**). For sequence recovery, a similar trend can be observed, except with AutoRotLib and MakeRotLib trading positions. The high performance of the “parent” approach is unsurprising, considering its adherence to default Rosetta parameters that were used to fit the Rosetta energy function. The decrease in its performance compared to default Rosetta is likely due to the use of UFF optimized geometry. The next best performing method is our method, FakeRotLib. However, we recognize that this comparison may be unrepresentative of more esoteric amino acid identities due to the inclusion of experimental torsional libraries in the RDKit conformational generator.^26^ Therefore, we have also included performance for a version of FakeRotLib which excludes these libraries, which caused the performance to drop between MakeRotLib and AutoRotLib. MakeRotLib, the traditional Rosetta tool to generate NCAA rotamers offers good performance, but not the best performance. AutoRotLib performs similarly to MakeRotLib, achieving neither superiority to FakeRotLib nor inferiority to the PDB rotamer methods. Finally, the PDB rotamer-based approaches perform the worst, but above the negative control of no rotamer information. Between these approaches, the RDKit-based approaches clearly outperform the BCL approach due to the inclusion of the aforementioned experimental torsion libraries.

**Figure 3.**
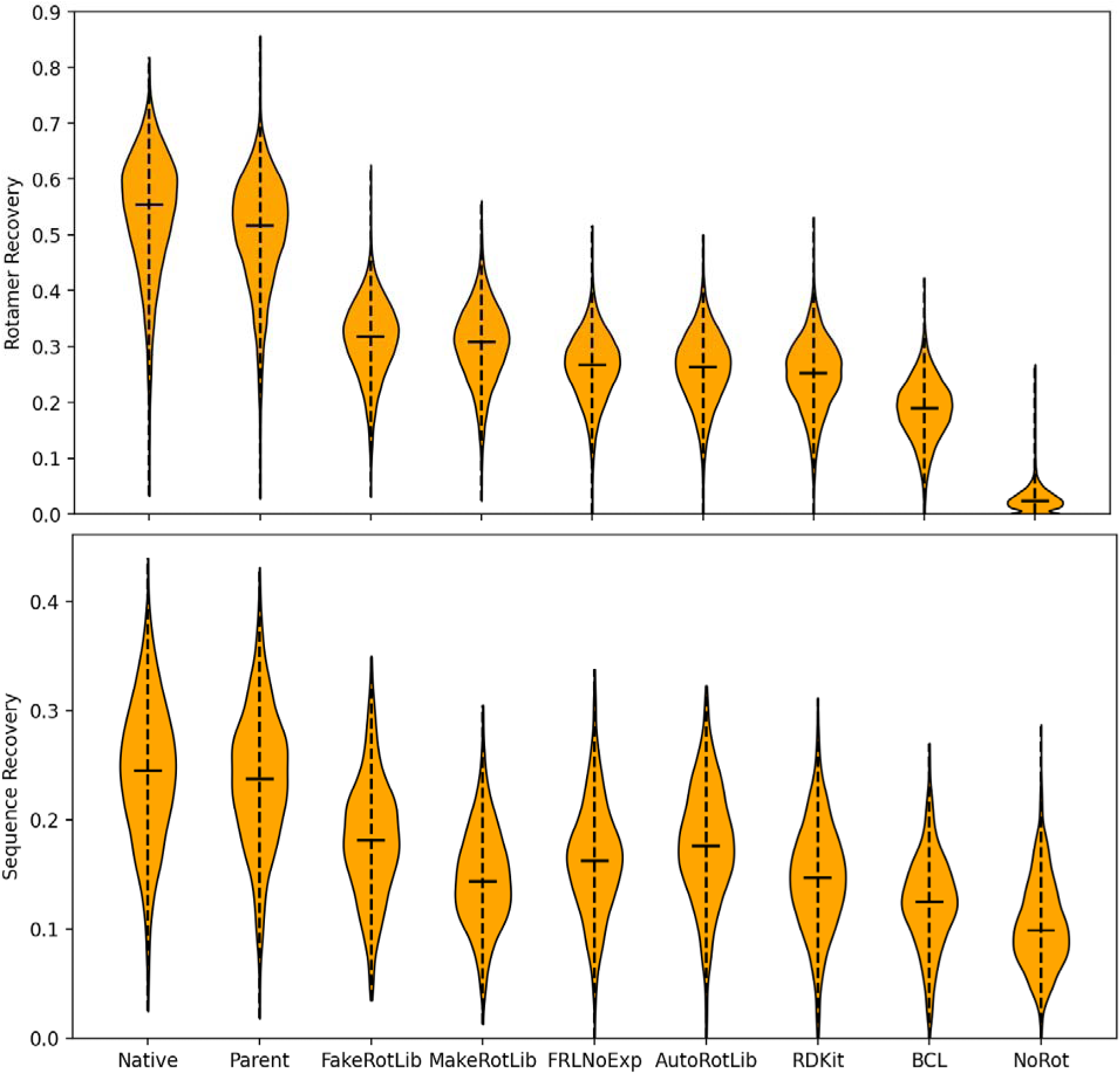
Rotamer recovery (top) and sequence recovery (bottom) performance of default Rosetta (Native), “Parent” rotamers (Parent), FakeRotLib, MakeRotLib, FakeRotLib without experimental torsions (FRLNoExp), AutoRotLib, PDB rotamers via RDKit, PDB rotamers via BCL, and no rotamer library (NoRot).

As a visualization of the rotamers being generated by these different approaches, we show the rotamer distribution of leucine as generated by each approach (**Figure 4**). For all methods except BCL and RDKit, the distributions were generated by a Metropolis-Hastings simulation; for the BCL and RDKit methods, we plot the dihedral distribution of an equal amount of PDB rotamers. The “native” distribution clearly replicates the original Dunbrack rotamer library,^27^ and as expected, the “parent” approach closely follows. Also expected is the diffuse nature of the distribution without provided rotamers, which allows almost all poses except those which cause internal steric clashing. Out of the remaining methods, FakeRotLib gives the best approximation of the native rotamer distribution, with only some problems of well standard deviation. AutoRotLib and FakeRotLib without experimental torsions both give faithful distribution approximations but have some clear smearing between wells or wide well standard deviations that potentially let through non-rotameric conformations. MakeRotLib’s distribution loosely resembles the native distribution but is clearly malformed. This is potentially due to our agnostic approach toward choosing initial centroid guesses for MakeRotLib. Finally, the PDB rotamer distributions hit the nine-well rotamer distribution, but the wells have contrasting issues: the RDKit distribution adheres too tightly to the experimental torsion library, resulting in tight wells around the “ideal” torsion angles, and the BCL wells are malformed and have clear connections between wells representative of non-rotameric sidechain poses.

**Figure 4.**
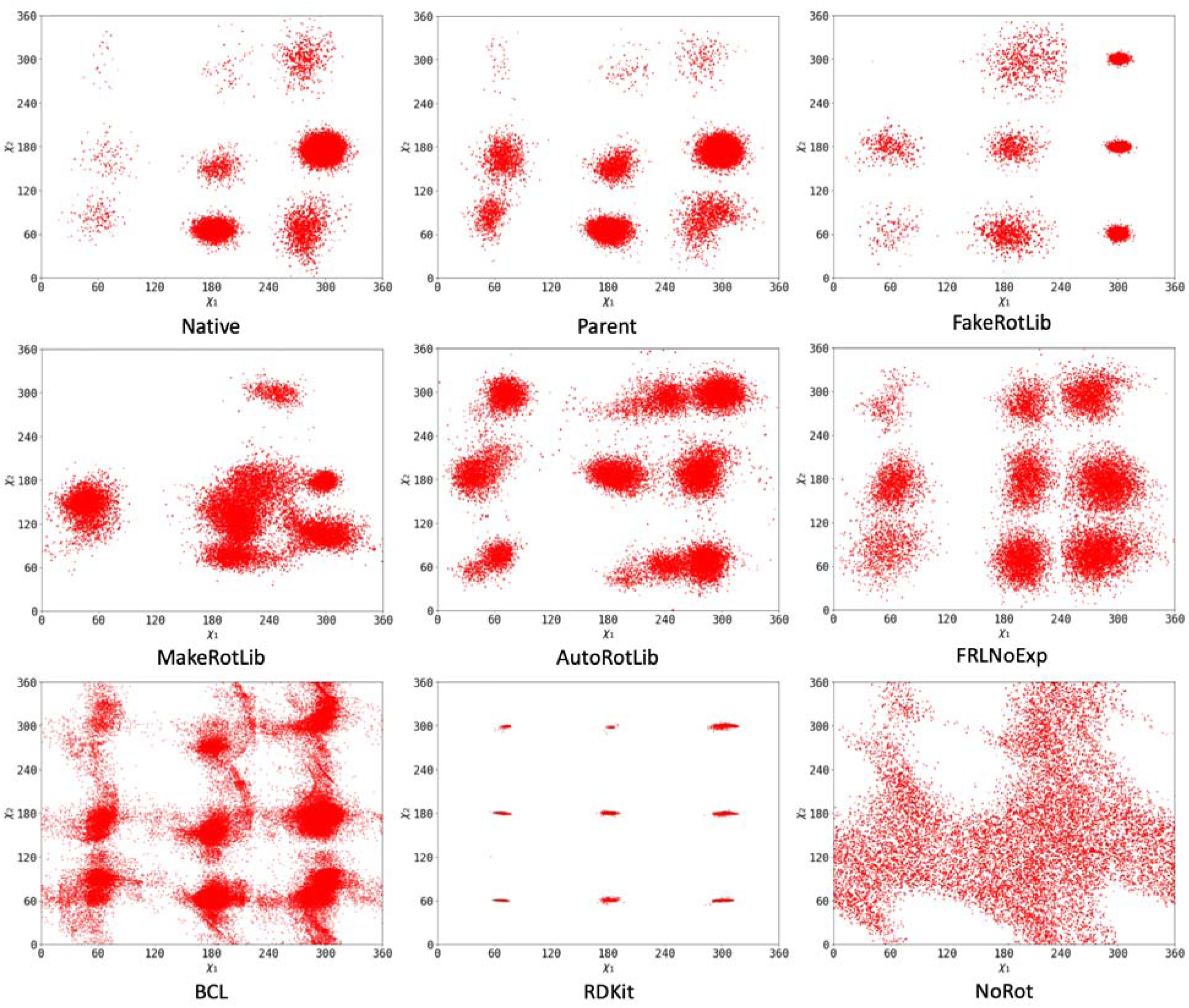
Rotamer distributions of leucine for every rotamer generation method: default Rosetta (Native), “parent” rotamers (Parent), FakeRotLib, MakeRotLib, AutoRotLib, FakeRotLib without experimental torsions (FRLNoExp), BCL PDB rotamers (BCL), RDKit PDB rotamers (RDKit), and no rotamers (NoRot).

It should be noted that the dihedral distributions of the PDB rotamers and the rotlib simulations exist in slightly different contexts; while both aim to show the dihedral distributions available to each given rotamer parameterization, the former is generated purely through small molecule conformer generation while the latter is the result of incorporating the rotamers into Rosetta. In addition, the number of rotamers shown in this plot is unrealistically high relative to the actual use case of the PDB rotamer sets; if PDB rotamers were used for actual design studies, these dihedral distributions would be much less populated. The reason for having so few PDB rotamers is due to Rosetta’s inefficient implementation of modeling involving these rotamer sets, particularly in its runtime scaling with increasing rotamer count. Even with the 100 rotamers per residue scheme we employed, some targets of the benchmark set would take days of runtime to finish a design run. While it is true that most Rosetta runs will not have every position be an NCAA as was benchmarked in this manuscript, this inefficiency is one which will hamper efficient design involving NCAAs in future studies.

## Discussion

As the results of this benchmark have demonstrated, the use of non-canonical amino acids in Rosetta remains a challenge. This is largely because the Rosetta energy function is built upon statistical distributions and performance optimizations based on experimental ground truth. However, such ground truth is inherently sparse in the case of non-canonical amino acids, as their chemical diversity extends far beyond what can be expected to be well-represented in databases. Therefore, given the current understanding of different parameterization methods available in Rosetta, one should try to adhere as closely as possible to default Rosetta parameters. For non-canonicals which are merely modifications of canonical amino acids, one should use the “parent” rotamer approach using default geometry for the substructure which matches the canonical and UFF geometry optimization on the portions which are divergent. However, for all other cases, we recommend using the FakeRotLib protocol which we have presented in this manuscript, generating a “rotlib” file for this protocol when the file format is supported (i.e., the amino acid is mono-substituted and has four or fewer chi angles). First of all, the performance using this parameterization scheme was shown to be superior to analogous methods in the above benchmarks. Secondly, the speed and convenience of this approach is much improved relative to existing tools: the entire protocol fits within one python script, demands only scikit-learn^28^ and RDKit^21^ be installed, and completes parameterizations of even highly flexible molecules within seconds. Finally, the automation of PDB rotamer library generation offered by FakeRotLib allows NCAA types previously unsupported by existing protocols to be parameterized, such as sidechains which conjugate with the backbone (e.g., proline derivatives) or highly flexible sidechains with more than four chi angles.

These recommendations are made given the current state of the default ref2015 Rosetta energy function,^29^ whose construction is exclusionary towards non-canonical amino acids (e.g. the reference energy is difficult to estimate properly for non-canonicals, ring closure applies only to prolines by default, sidechain geometries are hyperoptimized and difficult to replicate, terms such as the Ramachandran and amino acid probabilities are conditioned on amino acid type and become meaningless for non-canonicals, etc.). Therefore, if development is to continue in modeling non-canonical amino acids, serious consideration needs to be made with respect to how to fairly generate and evaluate conformations of non-canonicals.

## Methods

### Amino acid parameterization

#### Atom types and partial charges

We used three different methods for parameterizing atoms of each residue. The first is to use the atom types and partial charges automatically assigned by molfile_to_params_polymer.py. In this script, atom types are assigned based on atomic connectivity; partial charges are assigned to each atom as constants based on the atom type assignment and adjusted so that the sum of partial charges equals the formal charge. The second method is to use BCL to generate a partial charge file, and pass this partial charge file into molfile_to_params_polymer.py, thus leading to the same atom type assignments as the previous method but a more rigorous partial charge scheme. The final method is to use the atom types specified by the default Rosetta parameterization of each amino acid but reassign partial charges by drawing a sample the same size as the number of atoms in the residue from a uniform distribution over [-1,1] and recentering this sample so that the sum of all values is equal to the formal charge of the residue.

#### Sidechain geometry

Each canonical amino acid was first drawn in its zwitterionic form using Avogadro, and subsequently capped into a dipeptide form (i.e., the backbone was extended to the neighboring Cα on either side). For the “UFF” and “MMFF” geometries, this capping was performed through RDKit, and subsequently geometry optimized using the UFF and MMFF94 forcefields, respectively. The “QM” geometry was obtained through optimization of the UFF dipeptide via the Gaussian 16 software using the B3LYP method with a 6-31G(d,p) basis set. BCL geometry was accomplished using a custom applet which attaches the sidechain to an ideal glycine dipeptide backbone and generates a pose for the sidechain through the BCL conformer generator. However, this applet only properly functions for mono-substituted sidechains with at least one chi angle, i.e., proline, glycine, and alanine had to be treated separately. For these amino acids, the RDKit dipeptide was put into the BCL conformer generator, and the top-scored conformation was used as the BCL geometry.

#### Rotamers

Amino acids whose rotamers were derived through the “parent” method were assigned such that each canonical amino acid was assigned its own rotamer distribution. The “MakeRotLib” method refers to a previously published procedure in which residue starting torsions are iterated, each pose is minimized via a hybrid Rosetta/CHARMM energy function, and the resulting minimized structures are combined into rotamer wells. In running this protocol, an initial guess at the number of wells for each chi rotamer is required; we used 3 wells for the first chi angle of every residue and 6 wells for every subsequent chi angle. Also, since backbone conjugation causes errors in MakeRotLib, the “parent” method was used for proline in this method. For the “BCL” method, we passed the dipeptide into the BCL conformer generator, generated 1000 conformations, and kept the top 100 as PDB rotamers. The “RDKit” method was very similar, except that the conformation generation was carried out by RDKit, and the scoring was performed by UFF after removing the dipeptide extensions from the backbone (to ensure the rotameric energy of the sidechain and not the backbone was dominating the total energy).

#### FakeRotLib

“FakeRotLib” takes the set of PDB rotamers generated by the RDKit method and transforms them into Dunbrack-like rotamer wells, thus allowing for more flexible off-rotamer angle scoring. The protocol accomplishes this by fitting the dihedral angle distributions of the PDB rotamers via a Bayesian Gaussian Mixture Model (BGMM, a.k.a. an “Infinite Mixture Model”) as implemented by scikit-learn. Effectively, this model fits multivariate Gaussian peaks representative of the clusters in the dihedral distribution and draws the relative density of those peaks (as well as the optimal number of those peaks) through a Dirichlet process. In our implementation, we initialize the fitting with 10^*n*^ peaks (where *n* is the number of chi angles the sidechain has) each with an obligate diagonal covariance matrix (i.e., assuming dimension independence for the sake of simplicity).

Due to the incompatibility of Gaussian distributions with the modular number space of angular values, we fit the mixture model in cartesian space and convert the parameters of each Gaussian peak to angular values. First, each group of four atoms corresponding to each chi angle of the sidechain is superposed via Kabsch superposition against a reference frame constructed such that the central bond aligns with the Z axis and the first three atoms are aligned with the XZ plane. The XYZ position of the fourth atom given the superposition of the first three atoms against this reference frame is what is ultimately fit by the BGMM, resulting in 3*N-dimensional Gaussian peaks, where N is the number of chi angles. The cartesian means of each Gaussian peak are converted into dihedral space by calculating the dihedral angle between the mean XYZ position and the aforementioned reference frame. The standard deviation of each Gaussian peak is determined by finding a vector in the XY plane orthogonal to a vector pointing from the origin to the mean, finding where that vector intersects with the XY ellipse one standard deviation away from the mean, and calculating the absolute difference between the dihedral at that intersection point and the mean dihedral. With this new set of dihedral means and standard deviations, the 3*N-dimensional cartesian BGMM can be adapted into an N-dimensional dihedral BGMM.

Once the model has been transformed into dihedral space, all peaks with density less than 0.005 are discarded. The distribution of each chi angle is then fit independently as a one-dimensional BGMM, resulting in a number of bins for each chi angle corresponding to the number of Gaussian peaks with density above 0.005. Peaks in the full dihedral distribution are then recursively assigned a bin for each chi angle by finding the bin whose mean is closest to the peak’s mean in that dimension; if two peaks are given the same set of assignments, the peaks are merged via a weighted sum of means and covariances. Finally, the binned peaks are written to a “rotlib” file, where the same set of peaks are repeated for each phi and psi angle state of the backbone, thus making the rotamer set backbone independent. Note that because of limitations of the rotlib file format, a maximum of four chi angles are able to be represented; as a result, the fifth chi angle of arginine was removed for MakeRotLib and FakeRotLib. Also, to ensure well-formed dihedral distributions, ten times as many conformers are used in FakeRotLib compared to the RDKit PDB rotamers method.

### Benchmark Protocols

The rotamer recovery benchmark was carried out on a set of 12,357 non-redundant proteins from the CATH-S20 v4.3.0 dataset;^30^ a 1,000 protein subset of this set was used for the sequence recovery benchmarks. For both the rotamer and sequence recovery benchmarks, sidechain atoms were removed from these structures to ensure that the initial packing of the protein was not too proximal to the native structure. All residues of the protein are first mutated into their respective non-canonical forms. For the rotamer recovery benchmark, the residues are then packed by the PackRotamers mover. Rotamer recovery is then calculated from these structures as the percentage of residues whose chi angles are all within 20 degrees of the chi angles of the native structure. For the sequence recovery benchmark, we redesign the protein using the PackRotamers mover, restricting the set of residues for use in design to the specified non-canonical set. In this design, we set the reference energy of the Rosetta energy function to zero since non-canonical residues lack this energy term. Sequence recovery was then calculated as the number of residues whose respective canonical form matched the native residue.

To generate dihedral distribution plots for each of the parameterizations, we implemented a Metropolis Hastings simulation on a straight alanine 16-mer peptide using Rosetta Scripts.^31^ First, the eighth residue of this peptide is mutated into the non-canonical residue of interest, and this residue and its neighbors have its chi angles relaxed via FastRelax.^32–34^ Next, we run the Metropolis Hastings simulation for 10,000 steps of burn-in with two movements: a shear backbone movement and a sidechain rotation movement. We then record the next 100,000 simulation steps as a PDB trajectory, calculate the value of the chi angles of the mutated residue at each step using CPPTRAJ,^35^ and plot the resulting distribution.

## Author Information

### Corresponding Author

Dr. Jens Meiler; Institute for Drug Discovery, Institute for Computer Science, Wilhelm Ostwald Institute for Physical and Theoretical Chemistry, University Leipzig, Leipzig, Germany; Center for Scalable Data Analytics and Artificial Intelligence ScaDS.AI and School of Embedded Composite Artificial Intelligence SECAI, Dresden/Leipzig, Germany; Department of Chemistry, Department of Pharmacology, Center for Structural Biology, Institute of Chemical Biology, Center for Applied Artificial Intelligence in Protein Dynamics, Vanderbilt University, Nashville, Tennessee, United States of America. Address: 5114B MRB III, 465 21^st^ Ave S, Nashville, TN 37232.

### Author Contributions

E.W.B. was the primary author of the manuscript, including developing the FakeRotLib script, running benchmarks, generating figures, and writing the text. B.P.B. initially conceived the idea of using small molecule conformers for NCAA rotamers, created an early prototype of the FakeRotLib method, and contributed code for generating the rotamer density plots. J.M. oversaw the research and provided intellectual guidance. All authors assisted in and approve of the revision and preparation of this manuscript for publication.

## Acknowledgements

We would like to acknowledge Oanh Vu, who contributed significantly to the development of the molfile_to_params_polymer.py script and whose initial work on peptide modeling using NCAAs led to the creation of early FakeRotLib prototypes. We would also like to acknowledge Amanda Muyskens for suggesting the use of cartesian coordinates in clustering. OpenEye software licenses (needed for running AutoRotLib) were obtained via the OpenEye Academic License.

## Data and Software Availability

Protein structures and sequences are derived from publicly available data through the CATH database (https://www.cathdb.info/). FakeRotLib and other Rosetta packages are actively developed and maintained by the RosettaCommons (https://github.com/RosettaCommons/rosetta). The source code specifically used for this manuscript and raw data used to generate figures is made available at the author’s public fork of this repository (https://github.com/ewbell94/FakeRotLib).

